# HydraRNA: a hybrid architecture based full-length RNA language model

**DOI:** 10.1101/2025.03.06.641765

**Authors:** Guipeng Li, Feifei Jiang, Junhao Zhu, Huanhuan Cui, Zefeng Wang, Wei Chen

## Abstract

RNA, an essential component of the central dogma of molecular biology, plays versatile roles in all cellular processes. RNA large language models (LLMs) are emerging as powerful methods in RNA research to decipher its intricate network of function and regulation. However, previous RNA LLMs were based on the transformer model and pre-trained on short segment of non-coding RNAs, which limits their general usability. Here we present the first full-length RNA foundation model, HydraRNA, which is based on a hybrid architecture of bidirectional state space model and multi-head attention mechanism, and is pre-trained on a large amount of both protein-coding and non-coding RNAs. Despite being pre-trained with the fewest parameters and the least GPU resources, HydraRNA learns better RNA representations and outperforms the existing foundation models on a variety of downstream tasks, including RNA classification, prediction of RNA secondary structure, RBP binding sites, mRNA stability and translation efficiency. Furthermore, HydraRNA can accurately predict the effect of mutations and estimate the relative contributions of different mRNA regions to the RNA stability and translation. We anticipate that HydraRNA will enable dissecting the diverse properties of RNA, accelerating the research of RNA regulation and facilitating the optimal design of RNA therapeutics.

## Introduction

Ribonucleic acids (RNAs) can be categorized into two classes: protein-coding RNAs and non-coding RNAs. The former serves as carriers of genetic information in the central dogma of molecular biology, whereas the latter plays versatile roles in various important cellular processes^1^. The significance of RNA has been highlighted in numerous fields, from fundamental molecular biology and genetics to biotechnology and medicine^2,3^. For instance, the role of mRNA in the translation process has made it a focal point for RNA-based therapies, particularly in the context of mRNA vaccines for infectious diseases such as COVID-19^4,5^. Given the importance of RNA and the complexity of RNA biology, there has been increasing interest in the development of advanced computational methods for deciphering its intricate network of functions and regulations.

Large language models (LLMs), a form of artificial intelligence originating from natural language processing, have emerged as powerful methods in RNA research. By pre-training on vast amounts of RNA sequences and fine-tuning on experimental data, LLMs for RNA show promising performance in various RNA-related downstream tasks, such as predicting RNA functions and structures. RNA-FM^6^, the first RNA foundation model, was pre-trained on non-coding RNA segments from RNAcentral^7^ database and fine-tuned to solve multiple tasks including prediction of RNA secondary structures. RNAErnie^8^, which was built upon the Enhanced Representation through Knowledge Integration (ERNIE)^9^ framework, used a motif-aware pre-training strategy on RNAcentral database, and showed superiority in multiple tasks compared with other baselines^8^, including RNABERT^10^, RNA-MSM^11^ and RNA-FM. All these methods were essentially based on the transformer^12^ architecture. However, since attention mechanisms scale quadratically with sequence length^12^, these models limit their input size, which are often unable to handle full-length RNA sequence as a whole. Moreover, current models were pre-trained on truncated short segments of non-coding RNAs, which may hamper its power in full-length mRNA-related tasks. In addition to these general RNA LLMs, there are also specialized RNA LLMs, such as 3’UTRBERT^13^ and UTR-LM^14^, which focus only on the untranslated regions (UTRs) of the mRNA. These methods, by design, are not optimal for general tasks. Indeed, whether such specialized RNA LLMs have an advantage over general RNA foundation model even on UTR-related tasks is also controversial. Moreover, coding sequences are well-known to affect the mRNA translation^15,16^. Therefore, although predicting the effect of 5’UTR or 3’UTR on RNA properties, such as mRNA translation efficiency and stability, is valuable for designing sequences of mRNA therapeutics, the accurate prediction of the properties of mRNA in its full-length would be necessary for the further sequence optimization and is increasingly relevant with the rise of RNA-based therapies. To accelerate the AI-empowered research in RNA biology and medicine, there is a growing demand for a new RNA foundation model that can handle both non-coding and protein-coding RNA in its full-length .

To tackle these problems, we developed a new full-length RNA language model, HydraRNA.We applied HydraRNA on multiple RNA-relevant tasks, including RNA classification, prediction of RNA secondary structure, RBP binding sites, mRNA stability and translation efficiency. HydraRNA outperformed current RNA LLM models across all these downstream tasks. Furthermore, HydraRNA accurately predicted the effects of mutations and estimated the relative contribution of different parts of mRNA to the RNA stability and translation. Overall, we demonstrate HydraRNA as a valuable method for dissecting the diverse properties of RNA , which could further accelerate the AI-empowered research in RNA biology and medicine.

## Results

In the following sections, we first introduce the development of HydraRNA. Then, we present the experiment results for HydraRNA evaluation, which includes both unsupervised and supervised learning. Our analysis include evaluation not only at RNA segment level (5’UTR and 3’UTR), but also at the full-length RNA level. Unless otherwise specified, a simple multi-layer perceptron (MLP) predictor with only one hidden layer was used for each downstream task.

### Development of HydraRNA

HydraRNA is based on a hybrid architecture consisting of both Hydra^17^ and multi-head attention (MHA)^12^ layers (Figure 1a, also see Methods). It has 12 layers in total. Each layer contains a Hydra module except the 6th and 12th layer, which contains a MHA module. Hydra is a natural bidirectional extension of the structured state space model^17^ (Figure 1b). HydraRNA takes advantages of Hydra’s linear scalability of sequence length and high efficiency in in-context learning. MHA layer was inserted into the stack of Hydra layers to further improve the model quality (Figure 1a). Each layer has a hidden state dimension of 1024. HydraRNA was pre-trained on 28 million sequences of non-coding and protein-coding RNA from RNAcentral and NCBI databases (Figure 1c). A random span masking strategy, which is a generalization of the classical BERT-like masking strategy (see Methods), was used in the pre-training step. To speed up training, RNA sequences longer than 4096 nt were split into segments of length at most 4096 nt. About 90% of RNAs were pre-trained as full-length sequences without segmentation ( Supplementary figure 1). It took only 92 hours to pre-train HydraRNA on a server with 8 4090D GPU cards. The time and budget of training was significantly reduced comparing with previous RNA LLMs ( Supplementary table 1).

**Figure 1.**
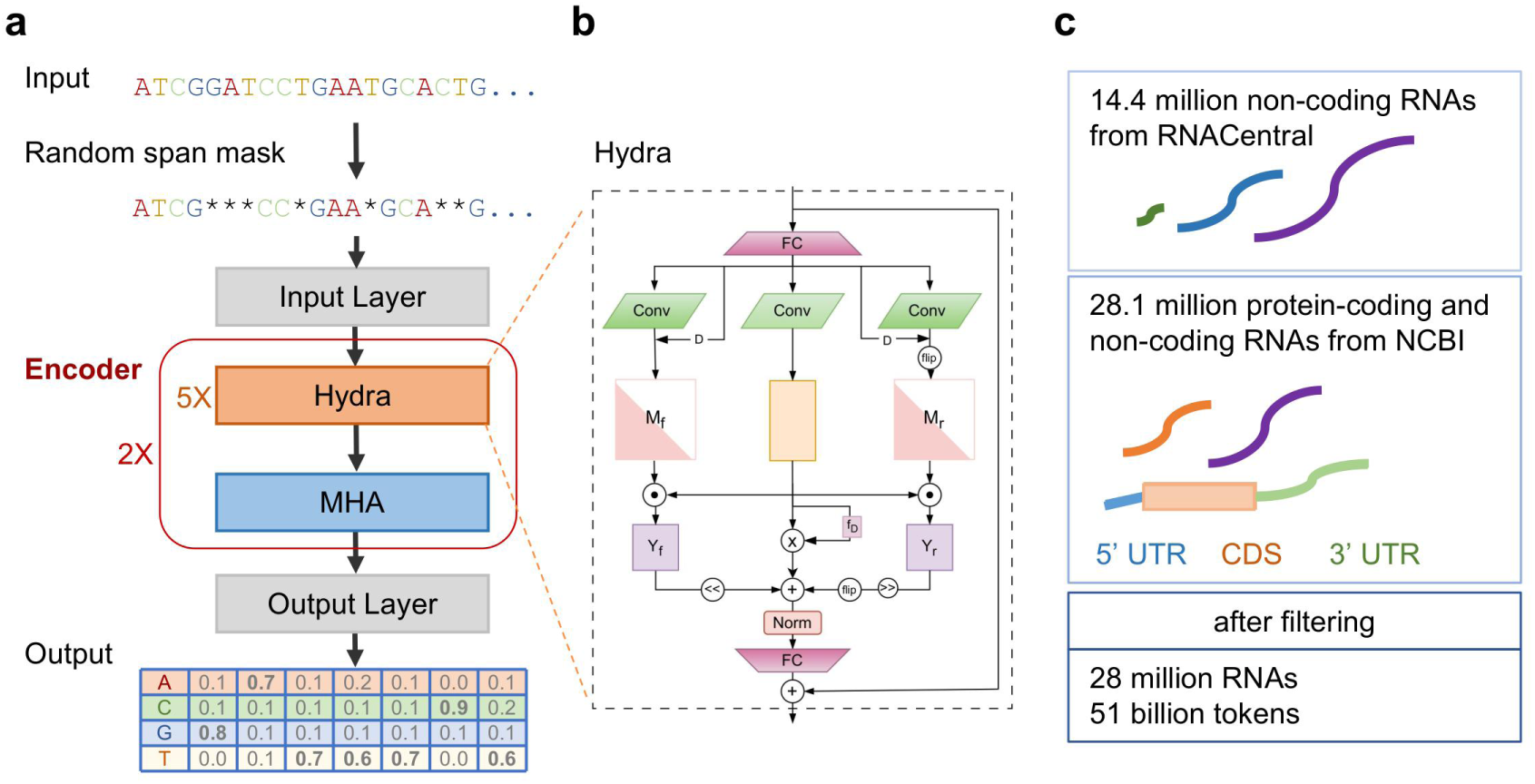
Overview of HydraRNA. **a**. HydraRNA employs a hybrid architecture with 12 layers. Each layer contains a Hydra module except the 6th and 12th layer, which contains a MHA module. **b**. Illustration of Hydra module. **c**. HydraRNA was pretrained on 28 million RNAs including both non-coding and protein-coding RNAs using masked language modeling.

### Visualization of the learned embedding

RNAs can be categorized into two types: protein-coding mRNAs and non-coding RNAs. It has been shown previously that the characteristics of various RNA types are partially captured within the embedding learned by the RNA language models^8^. To test the capability of the embedding to distinguish human protein-coding and non-coding RNA, we visualize the RNA sequences by employing t-SNE to the embedding in unsupervised setting and compare HydraRNA to two published RNA foundation model, RNA-FM and RNAErnie (Figure 2a). Since RNA-FM and RNAErnie were pre-trained only on non-coding RNA, they struggle to separate these two types of RNA. HydraRNA, in contrast, capture the intrinsic characteristics difference between these two types of RNA and the embedding exhibits a clearly separable clustering structure. We next test whether the embedding contain enough information to distinguish human protein-coding and non-coding RNAs in supervised classification task (Figure 2c). Again, HydraRNA achieve an AUC of 0.99, which outperforms the other RNA language models (0.88 and 0.71 for RNA-FM and RNAErnie, respectively). HydraRNA also outperforms the two models in terms of F1-score and accuracy (Figure 2c). The same trend holds for the mouse dataset (Supplementary figure 2). Considering only non-coding RNAs, HydraRNA shows comparable performance in distinguishing different classes of non-coding RNAs in both unsupervised (Figure 2b) and supervised setting (Figure 2d).

**Figure 2.**
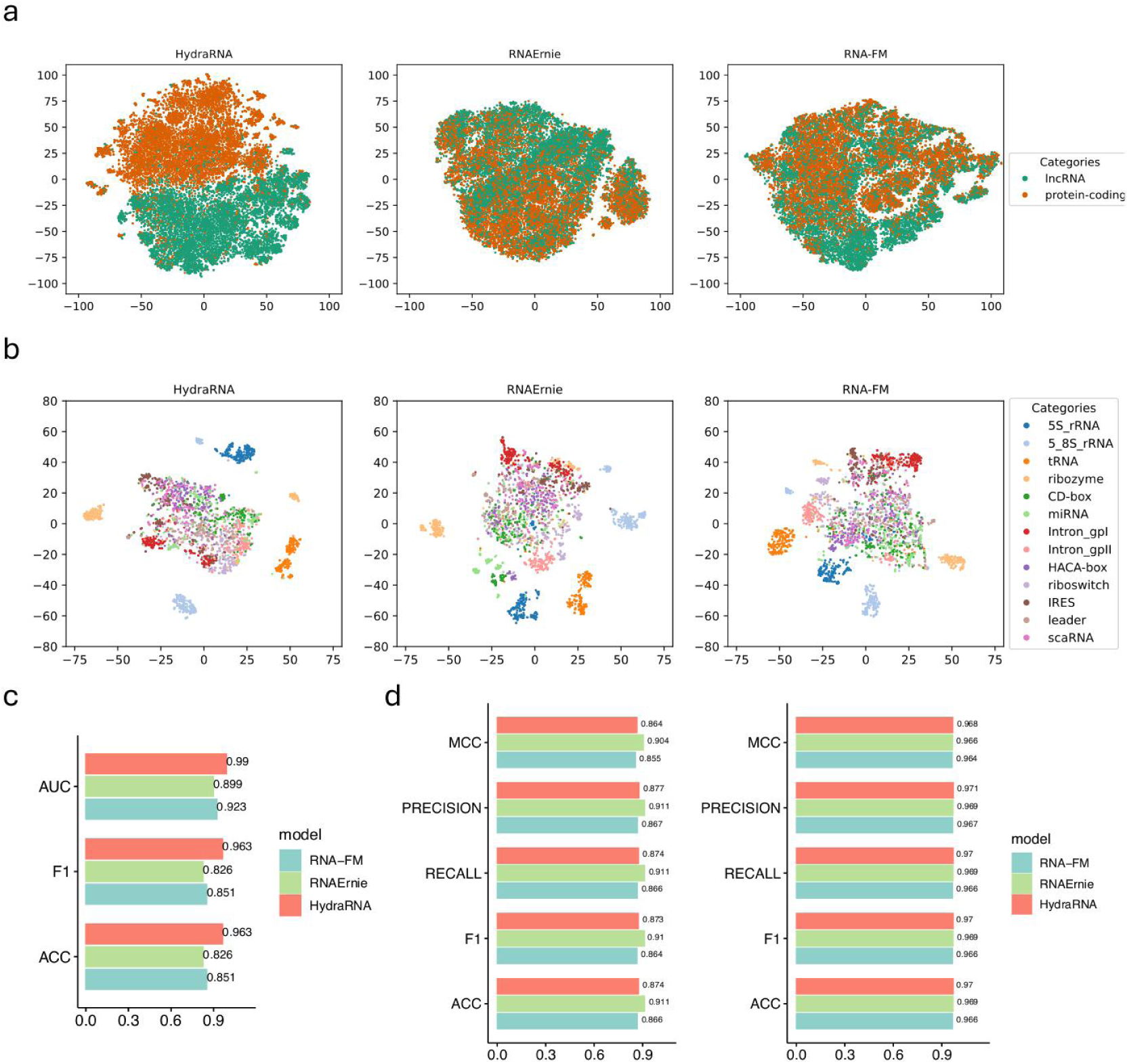
HydraRNA learned the intrinsic characteristics of RNA. **a**. The t-SNE 2D visualization of the embedding of protein-coding and lncRNA. **b**.The t-SNE 2D visualization of the embedding of 13 classes of ncRNA sequences. **c**. Model performance of binary classification for the lncRNA_H (human) dataset. **d**. Model performance of classification for the nRC dataset in frozen backbone setting (left) and fine-tuned setting (right).

### Prediction of RNA secondary structure

RNA secondary structure is vital for its stability and function. We utilized the dataset proposed in Singh et al.^18^ from bpRNA-1m database^19^ to benchmark the task of RNA secondary structure prediction. This dataset was split into three subsets: TR0 (10,814 structures) for training, TV0 (1,300 structures) for validation and TS0 (1,305 structures) for testing, and they are non-redundant on the sequence similarity level. The task is to classify each nucleotide pair as base-paired or unpaired. We fine-tuned HydraRNA on this binary classification task with binary cross-entropy loss, and compared HydraRNA to RNA-FM, RNAErnie and other popular methods specialized for secondary structure prediction, such as RNAfold^20^, SPOT-RNA^18^, UFold^21^ (Table 1) . The baseline values are all taken from cited literatures. HydraRNA achieves an impressive F1-score of 0.694 on the TS0 dataset, statistically higher than those of RNA-FM (0.676, p=0.000169) and RNAErnie (0.622, p<2.2e-16) (Figure 3a, Table 1).

**Figure 3.**
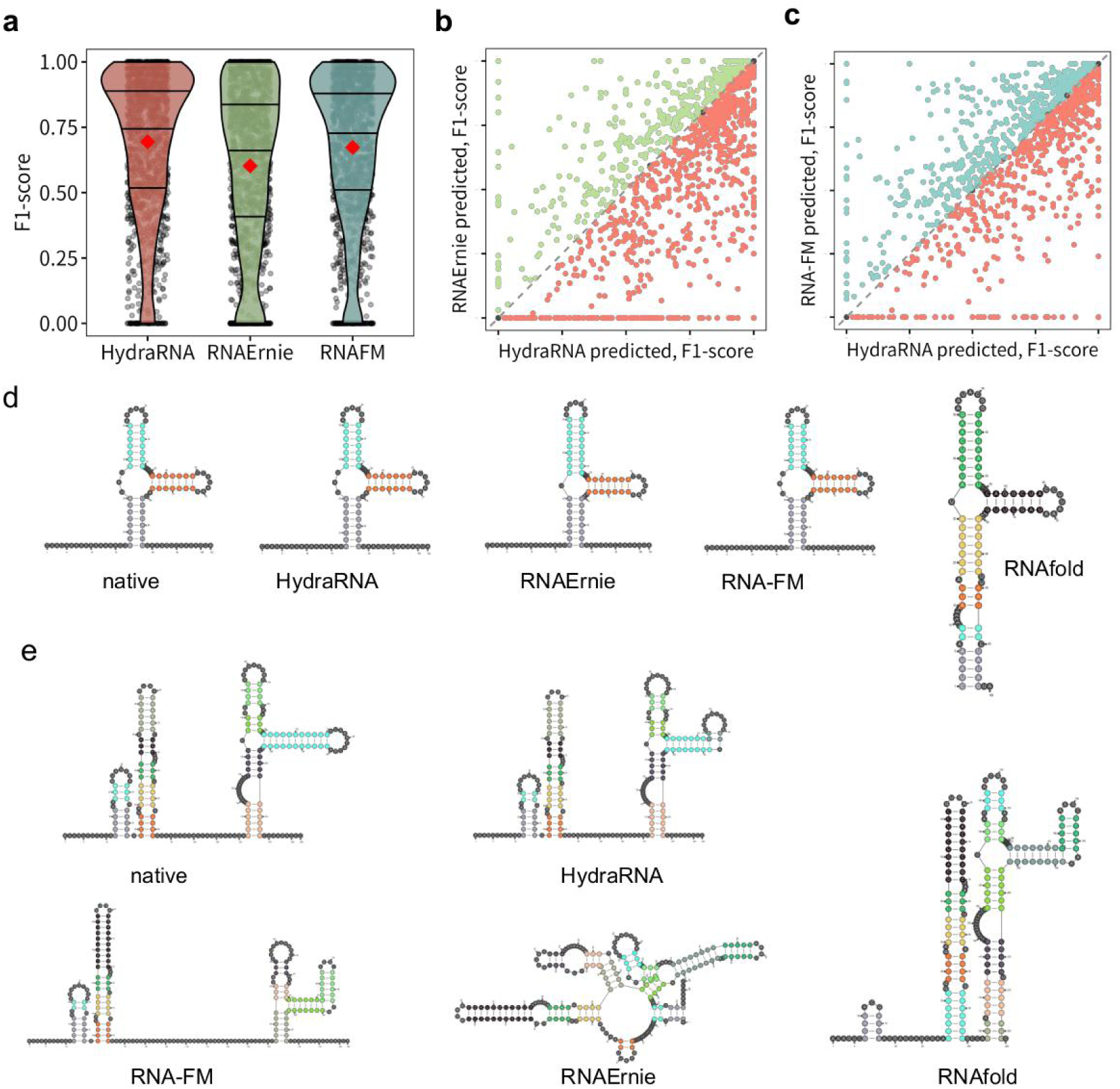
Performance of HydraRNA on RNA secondary structure prediction. **a**. Distribution of F1-score on the TS0 dataset between different models. **b**. Comparison of F1-score between HydraRNA and RNAErnie. **c**. Comparison of F1-score between HydraRNA and RNA-FM. **d**. Ground truth and prediction of the secondary structure of a purine riboswitch (bpRNA_RFAM_8591). **e**. Ground truth and prediction of the secondary structure of a group II intron (bpRNA_RFAM_38453).

**Table 1.**
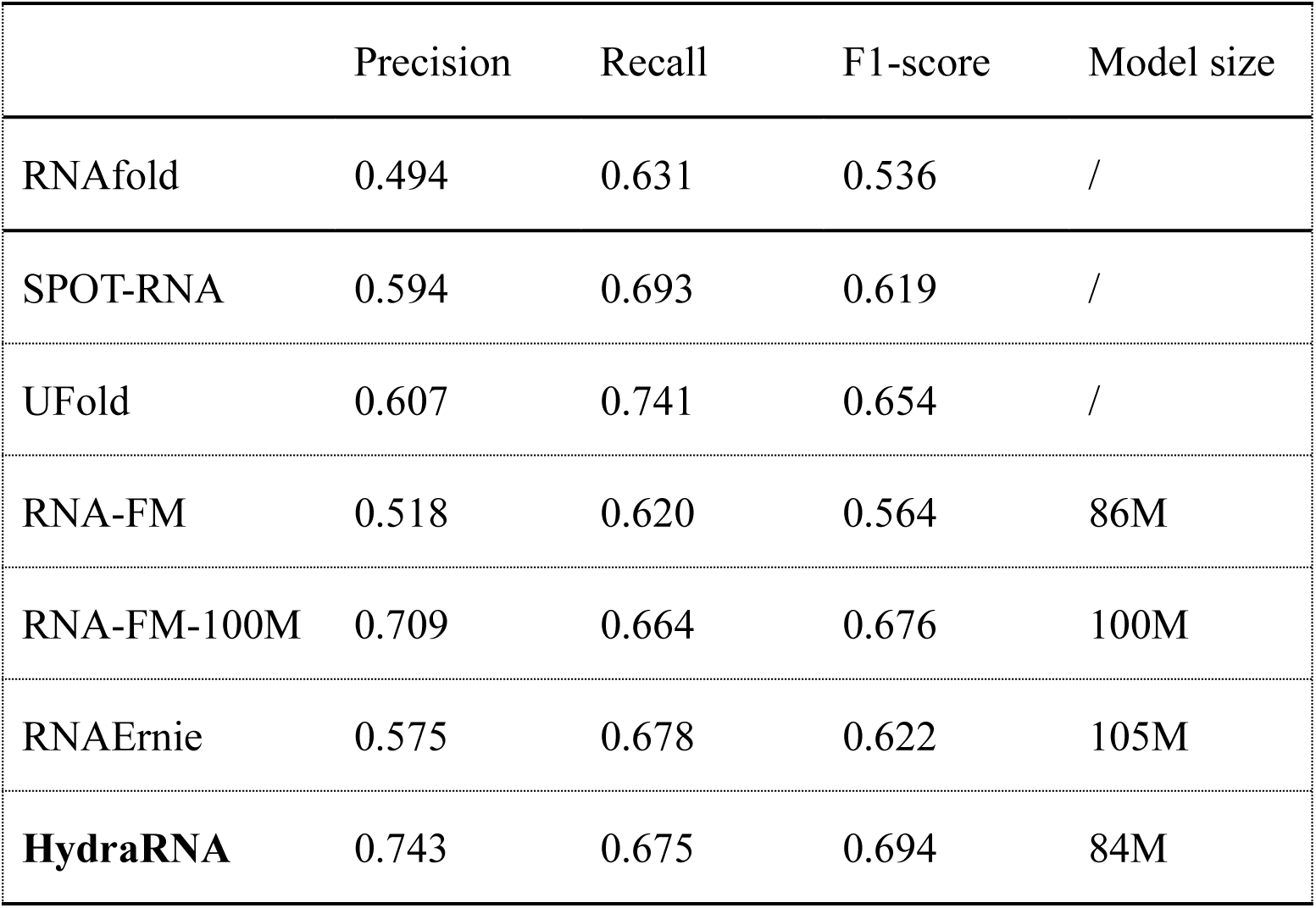
Performance of HydrRNA on RNA secondary structure prediction using TS0 dataset.

Among the 1,305 RNAs in TS0 test set, HydraRNA outperforms RNAErnie and RNA-FM-100M for 816 and 694 RNAs, respectively, while HydraRNA underperforms RNAErnie and RNA-FM-100M for 411 and 532 RNAs, respectively (Figure 3b, 3c). Whereas all the methods performed well for the relatively simple structure (Figure 3d), for the complex one, as shown in Figure 3e, the prediction from HydraRNA is much close to the native structure.

### Prediction of RNA-binding protein binding site

RNA-binding proteins (RBPs) typically recognize and bind to specific sequences or structures on the RNA transcripts to realize their regulatory functions. To benchmark the performance of HydraRNA on predicting the binding sites of RBPs, we used a dataset including 31 CLIP experiments on 19 RBPs curated by Yang et.al.^13^. For the benchmark study, we included RNA-FM, RNAErnie as well as 3URTBERT which previously showed state-of-the-art performance on this dataset^13^. As shown in Figure 4a, HydraRNA achieved the best performance compared to other methods (AUROC=0.866, AUPRC=0.682) (Figure 4a). Among these 31 CLIP experiments, HydraRNA outperforms RNAErnie, RNA-FM and 3UTRBERT in 25, 26, 22 experiments in terms of AUPRC metric, respectively (Figure 4b). Intriguingly, as shown in Figure 4b, the 31 CLIP experiments could be separated into two groups by the performance from all the four methods. For example, whereas all the four methods could predict well the binding sites of hnRNPC and eIF4AIII, the performances were all poor for NSUN2, AGO2 and IGF2BP1. We hypothesized that the prediction performance is dependent on the inherent sequence features determining RBP binding affinity. Some RBPs recognize and bind strongly to distinct sequence motifs, while the binding motif of other RBPs is more ambiguous. Conceivablely, it would be much easier for the models that have already learned the RNA sequence features by pre-training to predict the binding sites for the RBPs with strong and distinct binding motifs. Indeed, compared to those with satisfying prediction performance, the more challenging RBPs tend to have lower motif scores (Figure 4c, Supplementary table 2), indicating that RNA language models may contribute little to boost the prediction performance for these RBPs.

**Figure 4.**
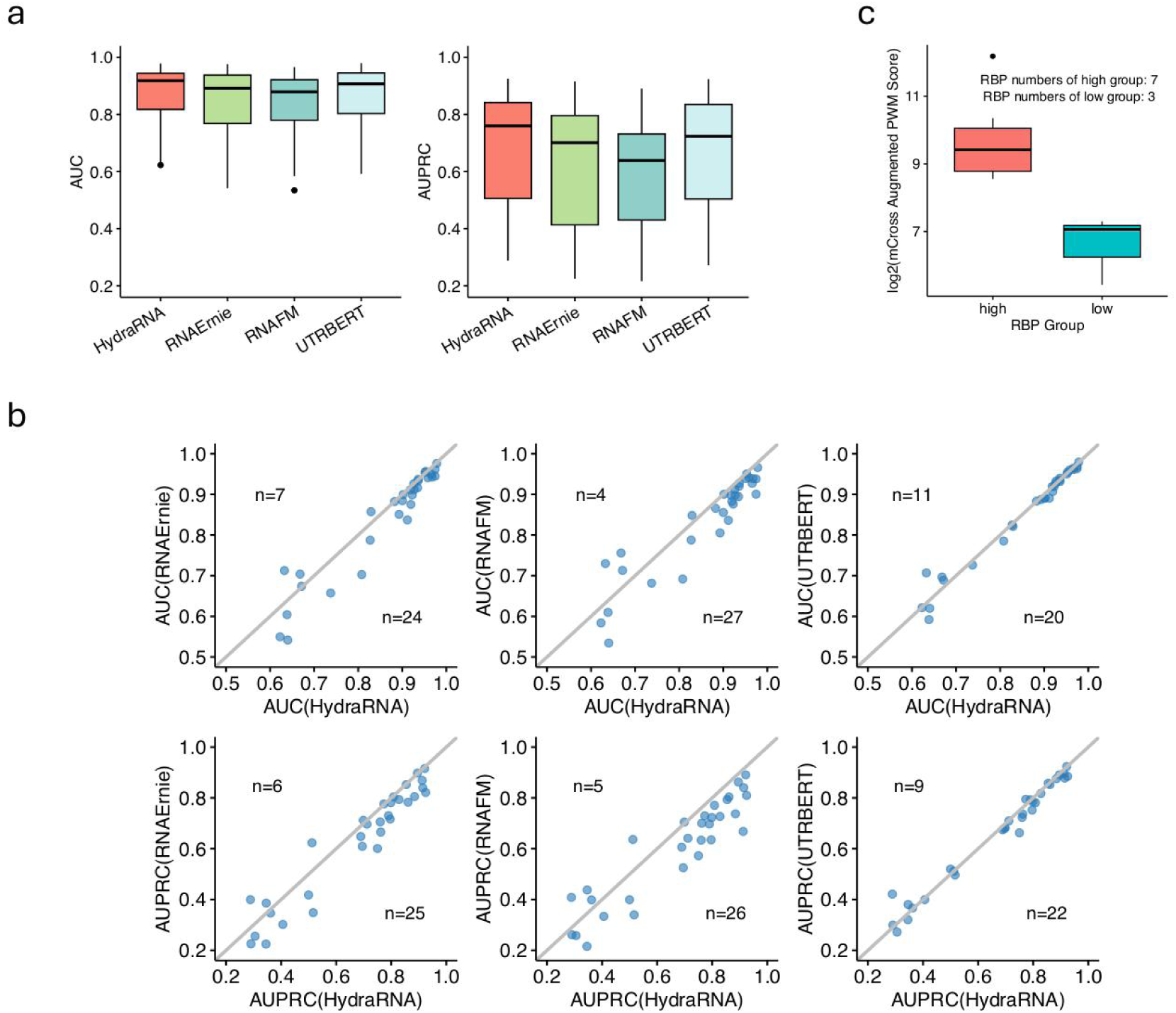
Performance of HydraRNA on predicting RBP binding sites. **a**. Box plot for comparison of AUC distribution.. **b**. Scatter plot for performance comparison. Each point represents one CLIP dataset. **c**. mCross augmented motif scores between two groups of RBP.

### Prediction of 5’UTR effect on ribosome loading

The 5’ untranslated region (5’ UTR) plays a critical role in regulating the translation from mRNA to proteins^22^. The mean ribosome loading (MRL), a widely used measure of mRNA translation efficiency, can be experimentally assessed through ribosome profiling. We evaluate the performance of our proposed model on a benchmark dataset^23^, which was also used in previous studies^6,14^. This dataset contains both random sequences and sequences from human 5’UTRs, with length ranging from 25 to 100 nt. We fine-tuned the HydraRNA on the 76,319 random 5’UTRs. Then we tested and compared HydraRNA with other baselines on 7,600 random and 7,600 human 5’UTRs, respectively. HydraRNA outperforms all the other methods in both tests, with Pearson R^2^ score of 0.914 in random test set (Figure 5a) and 0.85 in human test aset (Figure 5b). Notably, UTR-LM, a specialized 5’UTR language model, performs better than RNA-FM ,RNAErnie and Optimus on this task.

**Figure 5.**
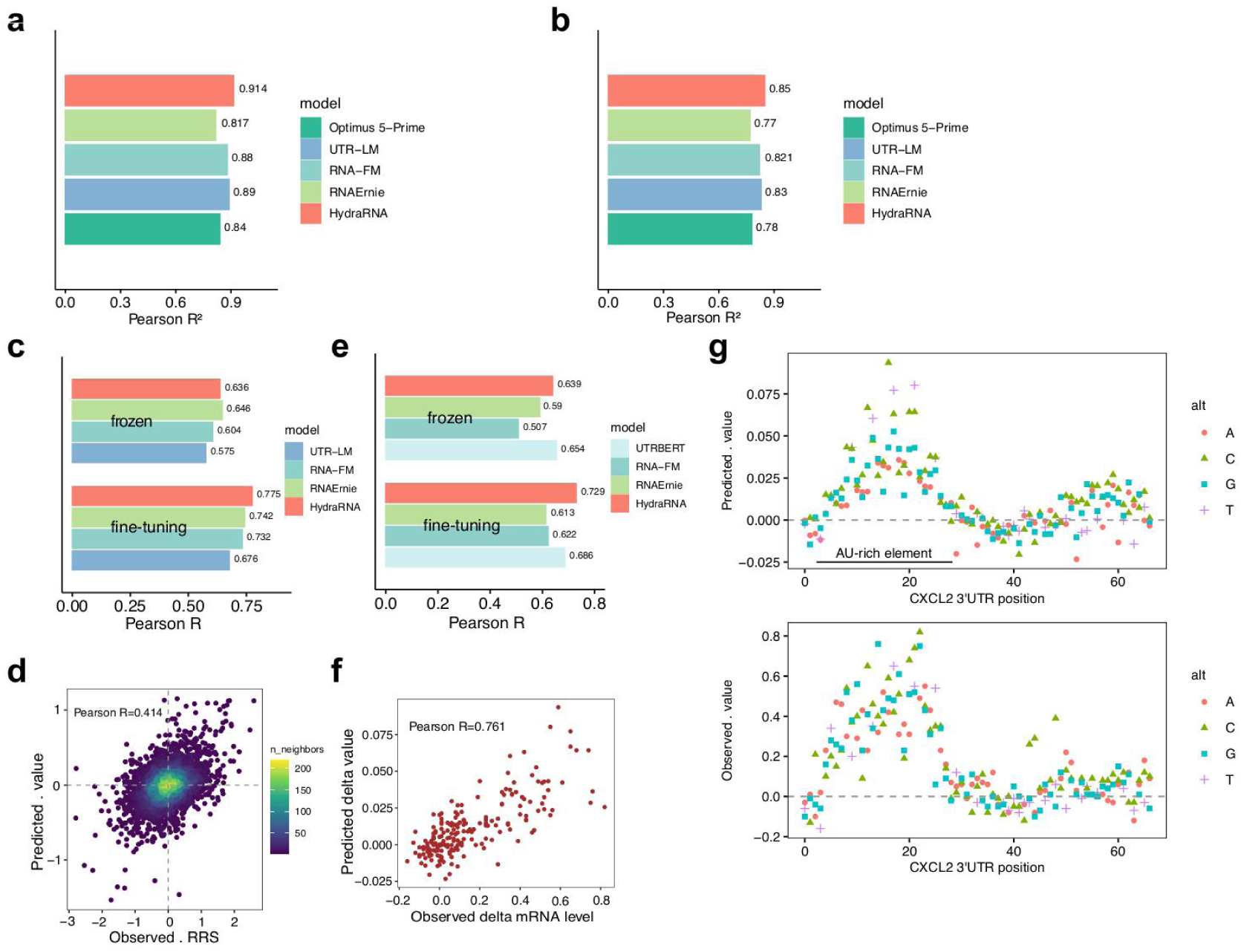
Performance of HydraRNA on predicting the effect of 5’UTRs on translation efficiency and 3’UTRs on RNA stability. **a**. the Pearson R^2^ value for the synthesized 5’UTRs dataset. **b**. the Pearson R^2^ value for the human 5’UTRs dataset. **c**. Performance comparison between models using frozen/all-parameters fine-tuning for the DART CleanCap dataset. **d**. Prediction of the impact of 5’UTRs mutation on translation efficiency. **e**. Performance comparison between models using frozen or all-parameters fine-tuning for the Beas2B dataset. **f-g**. Prediction of the impact of the 3’UTR mutations on RNA stability.

More recently, DART(direct analysis of ribosome targeting) has been developed to more directly measure the translation initiation rate^16^, which is believed to be the rate-limiting step for translational regulation. To evaluate the performance of HydraRNA for this type of data, we test HydraRNA on a recently published DART dataset, where ribosome recruitment scores were measured for more than 30,000 human 5’UTRs and their mutant variants^16^. As a result, HydraRNA achieved a Pearson correlation of 0.636 using frozen fine-tuning, which is comparable with RNAErnie (0.646) and higher than RNA-FM (0.604) and UTR-LM (0.575). After full parameter fine-tuning, the performance of HydraRNA was boosted to Pearson correlation of 0.775, which was higher than all the other baseline methods (Figure 5c). Notably, HydraRNA predict edthe effect of various mutations on ribosome recruitment reasonably well, with a Pearson correlation of 0.414 (Figure 5d). Assuringly, HydraRNA accurately predict the effect of most mutations with large effect size in the correct direction, indicating the potential use of HydraRNA in identifying the functional mutations from the whole genome or exome sequencing data from human patients.

### Prediction of 3’UTR effect on mRNA stability

Besides 5’UTRs, 3’ untranslated region (3’UTR) also contains cis-regulatory elements that control mRNA stability and translation ^24,25^. We then evaluated the performance of our model on a benchmark datasets where the effect of 3’UTR on mRNA stability were measured based on massively parallel reporter assays performed in human T lymphoblast Jurkat and human lung epithelial Beas2B cells^26,27^. This datasets were randomly split into training, validation and test set at 8:1:1 ratio. HydraRNA again outperforms both RNA-FM and RNAErnie with Pearson correlation of 0.729 in Beas2B (Figure 5c), which is higher than all the other baseline methods, including a specialized 3’UTR language model, 3’UTRBERT (R=0.686). This trend holds true for Jurkat dataset as well (Supplementary figure 3).We also tested the ability of HydraRNA in cross-celltype prediction (Supplementary figure 4). After fine-tuning in Beas2B dataset, the prediction performance in Jurkat is still high (Spearman R=0.57) but lower than within-celltype performance (Spearman R=0.66 for Beas2B, and Spearman R=0.75 for Jurkat). We further tested the ability of HydraRNA in predicting the effect of mutations on mRNA stability (Figure 5f). In the same study generating the benchmark dataset, the RNA stability was measured for a fast-UTR systematic mutagenesis library that included 201 single-base substitutions within a 67-nucleotide region of CXCL2 3’UTR ^26^. We removed all the sequences derived from CXCL2 genes from the training dataset, so the CXCL2 dataset could be treated as an independent test dataset, since the wild-type and mutant CXCL2 sequences were not used in the training process. As shown in Figure 5f, HydraRNA accurately predicted the effect of mutations in CXCL2 3’UTR on RNA stability with a Pearson correlation of 0.761 (Figure 5f). Notably, the mutations inside the AU-rich element substantially increased the RNA stability (Figure 5g), which is consistent with the well-known RNA distablizing effect of the AU-rich element ^28^.

### Prediction of mRNA translation and stability based on full-length mRNA sequences

Previous massive parallel reporter assay based screening usually used the same coding sequence (CDS), which often encodes fluorescent or luciferase proteins. Although the models trained on these datasets achieved impressive prediction accuracy on leave-out data, it has been shown that such models did not predict well the translation output of endogenous mRNAs^29^. The different CDS of endogenous mRNA likely contribute to the discrepancy in prediction performance. Then we evaluated the ability of HydraRNA in predicting the translation efficiency of the endogenous full-length mRNA, including 5’UTR, 3’UTR and CDS. This task is expected to be more challenging, especially for those models that were not designed for, thus could not learn the long context information of mRNA.

To benchmark, we collected a dataset where mRNA and protein abundance as well as their turnover were measured by parallel metabolic pulse labelling and translation rates were subsequently estimated based on kinetic modeling^30^. For this prediction task, HydraRNA achieved the highest Pearson R^2^. With the full-length mRNA sequence as input, HydraRNA could explained 41.3% of the variance in translation rate, while RNAErnie and RNA-FM only explained less then 18% (Figure 6a). To estimate the relative contribution of 5’UTR, CDS and 3’UTR to translation control, we further trained and tested HydraRNA on 5’UTR, CDS and 3’UTR sequences with the same data-split, respectively. Surprisingly, CDS alone could explained 40.4% of the variance in translation rate, which is almost the same as the full-length mRNA seqeunce (Figure 6a). 5’UTR alone explained 9.9% while 3’UTR alone only explained 4.5% of the variance. The same rank by contribution size (CDS>5’UTR>3’UTR) was revealed by other models as well (Figure 6a). For mRNA half-life prediction, HydraRNA also outperform RNA-FM and RNAErnie with a Pearson R^2^ of 0.32 (Figure 6b). The rank by contribution size for mRNA half-life from top to bottom is CDS (30.7%), 3’UTR (10%) and 5’UTR (0.7%). And again this pattern was also revealed by RNA-FM and RNAErnie (Figure 6b). As a control, in predicting the protein stablity, as expected, CDS explained much more variance than either 5’UTR or 3’UTR alone. Interesting CDS alone explained better than the full-length mRNA sequences, likely due to the noise attributed by UTR sequences (Figure 6c). To clarify whether this phenomenon is unique to this dataset, we examined it in another dataset which was generated by a modified pulsed-SILAC approach in mouse immune bone marrow-derived dendritic cells^31^. The prediction based on this dataset also revealed the same trend for translation rate (Figure 6d) and protein degradation rate (Figure 6e). Together, these results show that HydraRNA outperforms other RNA language models in long sequence related tasks and intriguingly, CDS contributes the most to mRNA translation rate as well as mRNA stability, compared with 5’UTR and 3’UTR. This emphasizes the important role of CDS in regulating mRNA fate, which needs to be considered in optimal design of mRNA therapeutics.

**Figure 6.**
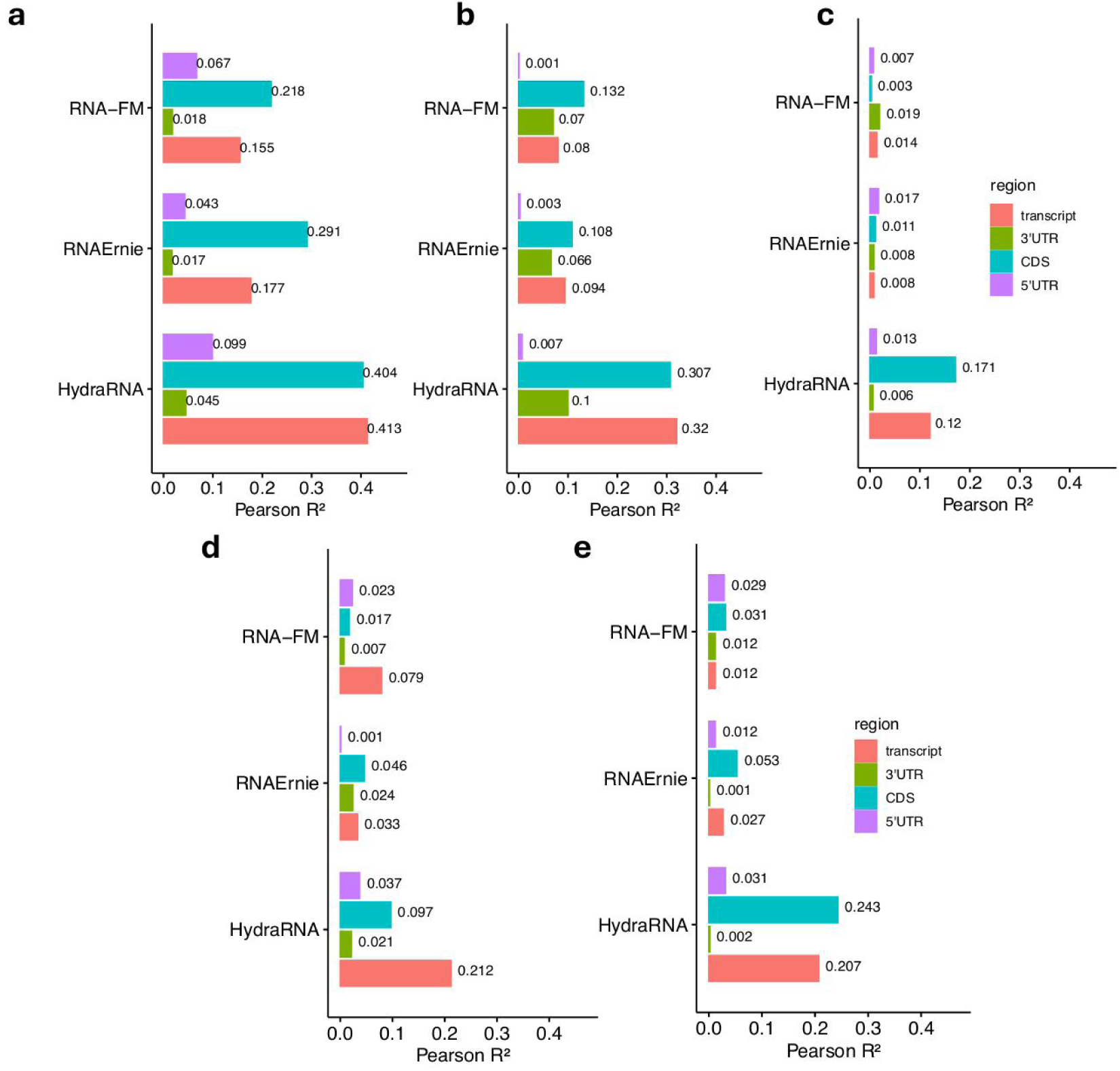
Prediction performance of HydraRNA on half-life and translation rate tasks. Prediction performance on translation rate (**a**), mRNA half-life (**b**), and protein half-life (**c**) from Schwanhäusser et.al.^30^. Prediction performance on translation rate (**d**) and protein degradation rate (**e**) from Jovanovic et.al.^31^.

## Discussion

Here, we developed a full-length RNA language model HydraRNA, which was built based on a hybrid architecture of Hydra and MHA. HydraRNA learned meaningful representations of RNA sequences by pre-training on a large corpus of both non-coding and protein-coding RNAs with a novel random span masking strategy. We demonstrated that HydraRNA achieved the state-of-the-art performance in diverse RNA-related tasks.

Notably, HydraRNA has a total of 84 million parameters. Although being trained with the least GPU resources, it outperforms other RNA LLMs for all the downstream tasks. This emphasizes the importance of model architecture, data processing and training strategy in a biological application of LLM. Two strategies incorporated in HydraRNA are essential for the higher performance. First, the unified random span masking strategy used in HydraRNA pre-training generalizes the classical BERT-like masking strategy to implicitly capture the deeper biological information underlying the consecutive region in the whole RNA transcript context, while it avoids explicitly using incomplete and biased motif priors (see Methods). Second, the lengths of mRNA and lncRNA often exceed thousands of nucleotide. Previous RNA foundation models used transformer architecture as the backbone, which is powerful in relatively short RNA and UTR learning, but with limited learning capability in long RNA context. Accordingly, these RNA foundation models were pre-trained on short segments of non-coding RNA sequences, which limited their power in diverse mRNA-related tasks. By integrating Hydra modules into its architecture, which scale linearly instead of quadratically with the length of input sequences, HydraRNA can handle long full-length RNAs efficiently and accurately. Taking advantage of this, we pre-trained HydraRNA with both non-coding and coding sequences. This together leads to a dramatic improvement over the other models in predicting mRNA translation and stability based on the full-length mRNA sequences, demonstrating HydraRNA as a powerful method to facilitate the study of RNA regulation as well as the optimal sequence design for mRNA therapeutics.

The performance of HydraRNA in different downstream tasks varies greatly, from R^2^ as high as 0.91 in predicting 5’UTR MRL to as low as 0.12 in estimating protein half-life. Even with the same kind of task, such as predicting the binding sties for different RBPs, the performance varies a lot. This is probably due to at least two reasons. First, the performance of any computational model relies on the quality of training data. Due to the nature of the biological problems as well as the complexity of experimental setups, some experiments could generate highly reproducible datasets whereas the other could not. Theoretically, the correlation between the biological replicates sets the ceiling for the model performance. Second, all the LLM models learn the sequence features by pre-training. They would not perform well for those tasks which are less determined by the RNA sequences, as exemplified in poor performance in predicting binding sites for the RBPs with ambiguous binding motifs.

It has been well recognized that mRNA translation and turnover could be regulated by cis-elements. Most of the known regulatory elements are located in the UTR regions ^32–34^. In particular, 5’UTR sequences are considered more important for modulating translation, whereas 3’ UTR sequences are more for mRNA stability. However, the relative contribution of these different regions in determining the translation and stability of mRNA has not yet been systematically quantified. Using HydraRNA with full-length mRNA sequence or only the sequence from isolated regions as input, we estimated for the first time the relative contributions of 5’UTR, 3’UTR and CDS in determining mRNA translation and stability. On the one hand, compared to 5’ UTR, 3’UTR sequence exerts more effect on mRNA stability, but less effect on mRNA translation, which is largely consistent with current knowledge. On the other hand, surprisingly, we revealed the dominate contributions of CDS on explaining the variance in translation rate as well as stability of endogenous mRNAs. We could not totally exclude that this is a technical artifact although two observations argue against this possibility. First, although dominant for mRNA translation, stability as well as protein half-life, the relative contributions of CDS are quantitatively very different. As a control, CDS explained much more variance than either UTR region, even marginally more than the the full-length mRNA sequences in predicting protein stability. This is expected given that the stability of protein would be largely determined by its protein sequences, which is solely based on the CDS. Second, the same trend was also found by the other two RNA LLM models, although they performed overall not as well as HydraRNA.

It should be also noted that there are limitations in this study. First, the HydraRNA model still includes two multi-head attention layers. This prohibits HydraRNA to handle much longer RNA sequences, such as 100K nucleotide in length. In this study, to balance the computation efficiency and full-length RNA context, RNA longer than 4K nucleotide were truncated in the pre-training step. But luckily, 88.3% and 99.0% of known mRNA are shorter than 4K and 10K nucleotide, so in most practical cases, 10K nucleotide would be sufficient. Second, RNA transcripts with high sequence identity were also excluded in the pre-training step. So the valuable evolutionary information encoded in these similar RNA transcripts between different species and isoforms were not fully utilized. Third, the current HydraRNA model is relatively much smaller than the LLM for protein sequences. Given that for any protein sequence, there are multiple mRNAs that could generate this same protein, the RNA sequence space is obviously much larger than the protein sequence space. While there are endeavors in scaling up the transformer-based RNA LLM^35 – 37^, it would be interesting to see the extent of gain in power when HydraRNA, a non-transformer-based method, scales up in the future study. Finally, in this study, HydraRNA was mainly used to predict the RNA sequence properties. Generative models could sample large amount of diverse RNA sequences from the unexplored RNA space. Integration of HydraRNA with generative models to efficiently generate and prioritize top RNA candidates with satisfied properties for wet-laboratory validation will be an exciting future direction in the field of RNA biotechnology and medicine.

## Methods

### HydraRNA architecture

We proposed a novel and hybrid architecture for RNA masked language modeling. The model is pretrained in a self-supervised learning manner via masked token reconstruction. Then it is partially or fully fine-tuned for a variety of downstream prediction tasks. The basic building blocks of HydraRNA include two types: Hydra and MHA. HydraRNA employs a 12-layer architecture and a hidden state dimension of 1024. Each layer contains a Hydra module except the 6^th^ and 12^th^ layer, which contains a MHA module. Hydra is a bidirectional structured state space model (SSM)^38^. Its strong performance in NLP and linear-time computational efficiency attract us to test whether Hydra is competitive in long RNA sequence modeling. Previous work suggest that a hybrid architecture with both SSM layers and attention layers could improve the model quality over that of a pure SSM (e.g., Mamba2) or a Transformer model^38^. Inspired by this result from NLP field, we inserted two MHA layers into the stack of Hydra architecture. FlashAttention^39^ was implemented in the MHA to speed up and reduce GPU memory usage.

For downstream task, we use the average embedding across the tokens of the sequence to represent the entire sequence. For fine-tuning the downstream tasks, unless otherwise specified, we freeze the pretraining model and only fine-tuning the MLP predictor which takes one dimensional embedding from RNA language model as input. The MLP predictor has only one hidden layer with 128 nodes. This configuration leads us to better quantify the quality of embedding from different RNA language models.

For RNA secondary structure prediction, inspired by RiNALMo^36^, we used a 12 layers 2D residual neural network (ResNet)^40^ as the prediction head, which is attached to the pre-trained HydraRNAmodel. An operation of outer concatenation is applied to the HydraRNA embedding to connect with the 2D ResNet.

### Random span masked language model

The masking strategy used in RNA-FM, RNABERT^10^, RNA-MSM^11^, LAMAR^41^, 3UTRBERT and UTR-LM was originally from BERT^42^, which is essentially at base-level. Specifically, 15% of the nucleobases are randomly selected within an RNA sequence. Among the selected tokens, 80% are masked, 10% are preserved without any alterations, and the remaining 10% are replaced with other nucleobases. Based on ERNIE, RNAErnie introduced a motif-aware multilevel masking strategy to pre-train the model. It maintained a proportion of 1:1:1 across the three different masking levels (base-level, subsequence-level and motif-level), with the training algorithm randomly selecting one strategy for each training session. Although this motif-aware masking strategy could incorporate biological priors into the training process, it may complicate the training process and biased to the known motif databases. We also noticed that another RNA foundation model, ERNIE-RNA^43^ also used ERNIE masking strategy during pretraining.

We proposed a novel unified masking strategy for masked language modelling, which is called random span masking. In this strategy, multiple span regions ( contiguous nucleobase sequences) that covering 15% of the nucleobases are randomly selected. Among the selected nucleobases, 80% are masked with <MASK> token, 10% are preserved without any alterations, and the remaining 10% are replaced with other nucleobases. The mean length of the span masks is set to be 4. This unified masking strategy generalize the classical BERT-like masking strategy to implicitly capture the deeper biological information underling the consecutive region in the whole RNA transcript context, while it also avoid explicitly using biased motif priors.

### Data for pretraining

RNA sequences from RNAcentral database were first filtered to include only RNAs with length between 18nt and 20,000nt, resulting in 32.5 million RNAs. Redundant and highly similar sequences were removed by using mmseqs^44^ with sequence identity cutoff at 0.8. 14.4 million RNAs were retained. 58.1 million RNA sequences from NCBI/RefSeq database were downloaded, which spread across 1,277 species. Redundant and highly similar RefSeq sequences were also removed by using mmseqs with sequence identity cutoff at 0.8, which results in 28.07 million RNAs. Then these 42.48 million filtered sequences from RNAcentral and NCBI were combined and processed by mmseq with sequence identity cutoff at 0.5. Finally 28.09 millions RNA sequences were used for pretraining, covering about 51 billion nucleotide tokens. The mean and median length of these RNA is 1811 and 1119 nt, respectively, with 89.7% shorter than 4096 nt. To speed up training, sequences longer than 4096 nt were split into segments of length at most 4096 nt. About 90% of full-length RNAs were pre-trained as a whole sequence without segmentation. Special token <S> and </S> were added at the start and end of each sequence during pretraining, respectively.

## Data for downstream tasks

The lncRNA_H, lncRNA_M and nRC datasets were obtained from the RNAErnie paper ( https://github.com/CatIIIIIIII/RNAErnie ). lncRNA datasets were employed for long-sequence binary classification (lncRNA/protein-coding RNA) using the human and mouse annotations. The nRC dataset was a multi-classification task, with sequences labelled into 13 RNA classes. Ning Wang et al.^8^ have pre-split the training and testing sets, which were also utilized in this research.

RBP binding sites datasets were downloaded from the UTRBERT paper (https://github.com/yangyn533/3UTRBERT). Yuning Yang et al.^13^ have collected data from 31 published CLIP experiments corresponding 19 RBPs. For each binding site prediction task, sequences of 101 nt were randomly split into training and validation sets at a 4:1 ratio. Additionally, an independent testing set containing 1000 sequences were used to assess the model’s generalization ability.

MRL datasets were downloaded from the Optimus 5-Prime paper^23^ (https://github.com/pjsample/human_5utr_modeling). The dataset includes 76,319 synthetic sequences for training, 7,600 for validation, and 7,600 endogenous sequences for testing.

DART CleanCap dataset and its corresponding variants dataset were acquired from Table S3 of Cole J.T. Lewis el al.’s paper^16^. Sequences were randomly divided into training, validation and testing sets with an 8:1:1 ratio. Data annotated in the variants dataset were removed during the training of the regression model.

Beas2B and Jurkat 3’UTR datasets were downloaded from https://github.com/david-a-siegel/AU-Rich-Elements. Sequences related to CXCL2 gene were removed. Sequences of 160 nt were used and randomly split into training, validation and test set at 8:1:1 ratio. CXCL2 mutation data of Beas2B cells were downloaded from https://www.nature.com/articles/nbt.2851#Sec17.

Full-length mRNA sequences with stability and translation label datasets were downloaded from Table S3 of Jovanovic et al.’s paper^31^ and Table S4 of Schwanhäusser et al.’s paper^30^. Data with missing values were excluded from subsequent analysis. Transcript annotations were downloaded from https://hgdownload.soe.ucsc.edu/goldenPath/mm9/bigZips/genes/mm9.refGene.gtf.gz, and the genome sequence was obtained from https://hgdownload.soe.ucsc.edu/goldenPath/mm9/bigZips/refMrna.fa.gz. Sequences from each region (5’UTR, CDS and 3’UTR) were extracted using bedtools^45^. Data lacking annotated 5’UTRs or 3’UTRs were removed. The remaining sequences were randomly divided into training, validation and testing sets with an 8:1:1 ratio.

### Hyperparameters and configurations

Fairseq^46^ framework was used for pretraining the HydraRNA model. In the pretraining stage, dynamic batch size was used. In each batch, max tokens per device was set to 40K. We utilized the Adam^47^ optimizer, which was regulated by a cosine learning-rate schedule involving anneal warm-up and decay. The initial learning rate was set at 1×10^-7^, with a maximum learning rate of 2×10^-4^. The learning-rate scheduler was designed to warm up during the first 10,000 steps. Gradients update frequency were set to 10. The model was pre-trained on eight Nvidia 4090D 24GB graphics processing units for around 90 hours.

## Data availability

All data used in this study were from publicly available datasets (see above).

## Code availability

HydraRNA can be accessed at HydraRNA Github repository ( https://github.com/GuipengLi/HydraRNA ), including the model weights and relevant source code.

## Acknowledgments

This work was supported by the National Key R&D Program of China (Grant No. 2021YFF1201000 and 2022YFC3400400 to W.C.), the Shenzhen Science and Technology Program (Grant No. KQTD20180411143432337 to W.C.) and the Shenzhen Key Laboratory of Gene Regulation and Systems Biology (Grant No. ZDSYS20200811144002008 to W.C.).

## Author contributions

G.L. and W.C. conceived the study. G.L. designed the method. F.J. and G.L. performed all the experiments except RNA secondary structure prediction. G.L. performed the experiments of RNA secondary structure prediction with the help of J.Z. G.L., F.J., H.C. Z.W. and W.C. discussed the results. G.L. and W.C. wrote the manuscript with the input from F.J, J.Z. H.C.. W.C. and G.L. coordinated and supervised the study.

## References

1. Sharp, P. A. The Centrality of RNA. Cell 136, 577–580 (2009).

2. Licatalosi, D. D. & Darnell, R. B. RNA processing and its regulation: global insights into biological networks. Nat Rev Genet 11, 75–87 (2010).

3. Parhiz, H., Atochina-Vasserman, E. N. & Weissman, D. mRNA-based therapeutics: looking beyond COVID-19 vaccines. The Lancet 403, 1192–1204 (2024).

4. Zeng, C. et al. Leveraging mRNA Sequences and Nanoparticles to Deliver SARS- CoV-2 Antigens In Vivo. Advanced Materials 32, 2004452 (2020).

5. Nance, K. D. & Meier, J. L. Modifications in an Emergency: The Role of N1-Methylpseudouridine in COVID-19 Vaccines. ACS Central Science (2021) doi:10.1021/acscentsci.1c00197.

6. Chen, J. et al. Interpretable RNA Foundation Model from Unannotated Data for Highly Accurate RNA Structure and Function Predictions. Preprint at http://arxiv.org/abs/2204.00300 (2022).

7. RNAcentral Consortium et al. RNAcentral 2021: secondary structure integration, improved sequence search and new member databases. Nucleic Acids Research 49, D212–D220 (2021).

8. Wang, N. et al. Multi-purpose RNA language modelling with motif-aware pretraining and type-guided fine-tuning. Nat Mach Intell 6, 548–557 (2024).

9. Sun, Y. et al. ERNIE 2.0: A Continual Pre-Training Framework for Language Understanding. AAAI 34, 8968–8975 (2020).

10. Akiyama, M. & Sakakibara, Y. Informative RNA base embedding for RNA structural alignment and clustering by deep representation learning. NAR Genomics and Bioinformatics 4, lqac012 (2022).

11. Zhang, Y. et al. Multiple sequence alignment-based RNA language model and its application to structural inference. Nucleic Acids Research 52, e3–e3 (2024).

12. Vaswani, A. et al. Attention Is All You Need. Preprint at http://arxiv.org/abs/1706.03762 (2017).

13. Yang, Y. et al. Deciphering 3’UTR Mediated Gene Regulation Using Interpretable Deep Representation Learning. Advanced Science 2407013 (2024) doi:10.1002/advs.202407013.

14. Chu, Y. et al. A 5 ′ UTR language model for decoding untranslated regions of mRNA and function predictions. Nat Mach Intell 6, 449–460 (2024).

15. Mauger, D. M. et al. mRNA structure regulates protein expression through changes in functional half-life. Proc. Natl. Acad. Sci. U.S.A. 116, 24075–24083 (2019).

16. Lewis, C. J. T. Quantitative profiling of human translation initiation reveals elements that potently regulate endogenous and therapeutically modified mRNAs. Molecular Cell (2024).

17. Hwang, S., Lahoti, A., Dao, T. & Gu, A. Hydra: Bidirectional State Space Models Through Generalized Matrix Mixers. Preprint at http://arxiv.org/abs/2407.09941 (2024).

18. Singh, J., Hanson, J., Paliwal, K. & Zhou, Y. RNA secondary structure prediction using an ensemble of two-dimensional deep neural networks and transfer learning. Nat Commun 10, 5407 (2019).

19. Danaee, P. et al. bpRNA: large-scale automated annotation and analysis of RNA secondary structure. Nucleic Acids Research 46, 5381–5394 (2018).

20. Lorenz, R. et al. ViennaRNA Package 2.0. Algorithms Mol Biol 6, 26 (2011).

21. Fu, L. et al. UFold: fast and accurate RNA secondary structure prediction with deep learning. Nucleic Acids Research 50, e14–e14 (2022).

22. Leppek, K., Das, R. & Barna, M. Functional 5 ′ UTR mRNA structures in eukaryotic translation regulation and how to find them. Nat Rev Mol Cell Biol 19, 158–174 (2018).

23. Sample, P. J. et al. Human 5′ UTR design and variant effect prediction from a massively parallel translation assay. Nat Biotechnol 37, 803–809 (2019).

24. Kuersten, S. & Goodwin, E. B. The power of the 3′ UTR: translational control and development. Nat Rev Genet 4, 626–637 (2003).

25. Bartel, D. P. MicroRNAs: Target Recognition and Regulatory Functions. Cell 136, 215–233 (2009).

26. Zhao, W. et al. Massively parallel functional annotation of 3 ′ untranslated regions. Nat Biotechnol 32, 387–391 (2014).

27. Siegel, D. A., Le Tonqueze, O., Biton, A., Zaitlen, N. & Erle, D. J. Massively parallel analysis of human 3 ′ UTRs reveals that AU-rich element length and registration predict mRNA destabilization. G3 Genes|Genomes|Genetics **12**, jkab404 (2022).

28. Barreau, C. AU-rich elements and associated factors: are there unifying principles? Nucleic Acids Research 33, 7138–7150 (2005).

29. Schlusser, N., González, A., Pandey, M. & Zavolan, M. Current limitations in predicting mRNA translation with deep learning models. Genome Biol 25, 227 (2024).

30. Schwanhäusser, B. et al. Global quantification of mammalian gene expression control. Nature 473, 337–342 (2011).

31. Jovanovic, M. et al. Dynamic profiling of the protein life cycle in response to pathogens. Science 347, 1259038 (2015).

32. Sonenberg, N. & Hinnebusch, A. G. Regulation of Translation Initiation in Eukaryotes: Mechanisms and Biological Targets. Cell 136, 731–745 (2009).

33. Shalgi, R. et al. Widespread Regulation of Translation by Elongation Pausing in Heat Shock. Molecular Cell 49, 439–452 (2013).

34. Jackson, R. J., Hellen, C. U. T. & Pestova, T. V. The mechanism of eukaryotic translation initiation and principles of its regulation. Nat Rev Mol Cell Biol 11, 113–127 (2010).

35. Wang, X. et al. UNI-RNA: UNIVERSAL PRE-TRAINED MODELS REVOLUTIONIZE RNA RESEARCH. Preprint at 10.1101/2023.07.11.548588 (2023).

36. Penić, R. J., Vlašić, T., Huber, R. G., Wan, Y. & Šikić, M. RiNALMo: General-Purpose RNA Language Models Can Generalize Well on Structure Prediction Tasks. Preprint at http://arxiv.org/abs/2403.00043 (2024).

37. Zou, S. et al. A Large-Scale Foundation Model for RNA Function and Structure Prediction. Preprint at 10.1101/2024.11.28.625345 (2024).

38. Dao, T. & Gu, A. Transformers are SSMs: Generalized Models and Efficient Algorithms Through Structured State Space Duality. Preprint at http://arxiv.org/abs/2405.21060 (2024).

39. Dao, T., Fu, D. Y., Ermon, S., Rudra, A. & Ré, C. FlashAttention: Fast and Memory-Efficient Exact Attention with IO-Awareness. Preprint at http://arxiv.org/abs/2205.14135 (2022).

40. He, K., Zhang, X., Ren, S. & Sun, J. Deep Residual Learning for Image Recognition. in 2016 IEEE Conference on Computer Vision and Pattern Recognition (CVPR) 770–778 (2016). doi:10.1109/CVPR.2016.90.

41. Zhou, H. et al. Deciphering RNA regulation with a foundation language model. Preprint at 10.1101/2024.10.12.617732 (2024).

42. Devlin, J., Chang, M.-W., Lee, K. & Toutanova, K. BERT: Pre-training of Deep Bidirectional Transformers for Language Understanding. Preprint at http://arxiv.org/abs/1810.04805 (2019).

43. Yin, W., et al. ERNIE-RNA: An RNA Language Model with Structure- enhanced Representations.

44. Hauser, M., Steinegger, M. & Söding, J. MMseqs software suite for fast and deep clustering and searching of large protein sequence sets. Bioinformatics 32, 1323– 1330 (2016).

45. Quinlan, A. R. & Hall, I. M. BEDTools: a flexible suite of utilities for comparing genomic features. Bioinformatics 26, 841–842 (2010).

46. 46. Ott, M., et al. fairseq: A Fast, Extensible Toolkit for Sequence Modeling. Preprint at 10.48550/arXiv.1904.01038 (2019).

47. Kingma, D. P. & Ba, J. Adam: A Method for Stochastic Optimization. Preprint at 10.48550/arXiv.1412.6980 (2017).

48. Cao, J. et al. High-throughput 5 ′ UTR engineering for enhanced protein production in non-viral gene therapies. Nat Commun 12, 4138 (2021).

