## Supplementary for "HydraRNA: a hybrid architecture based full-length RNA language model"

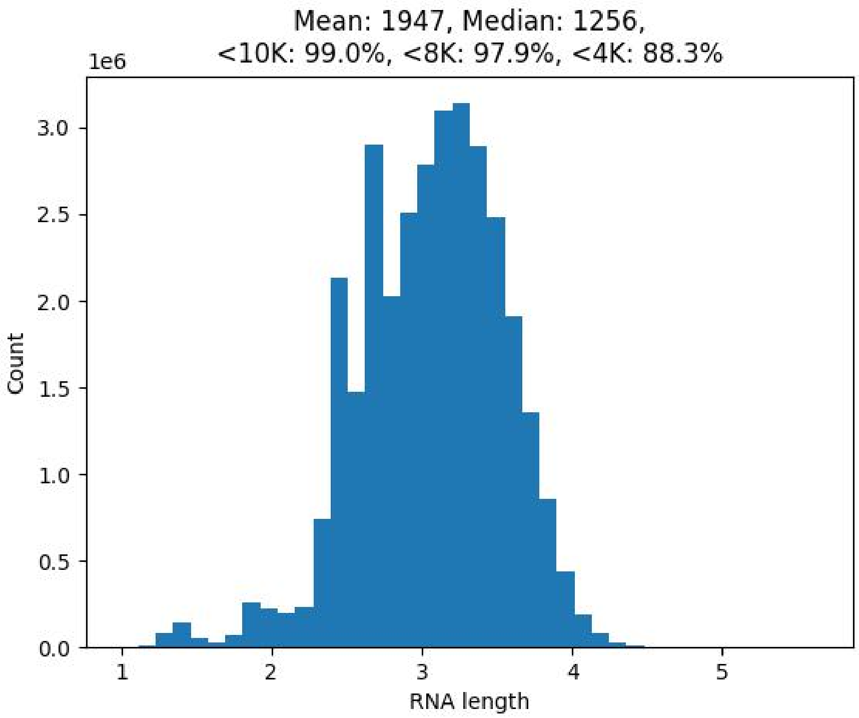


Figure S1. RNA length distribution. 88.3% of RNAs are shorter than 4096 nt.

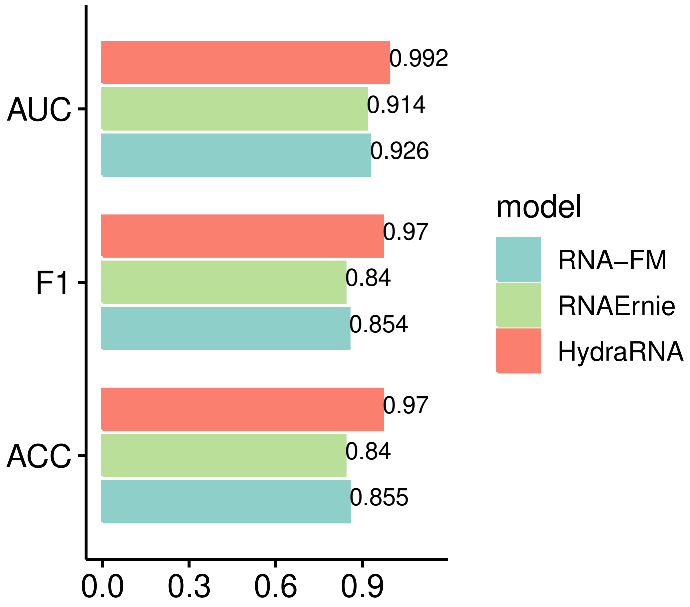


Figure S2. Model performance of binary classification for the lncRNA_M (mouse) dataset.

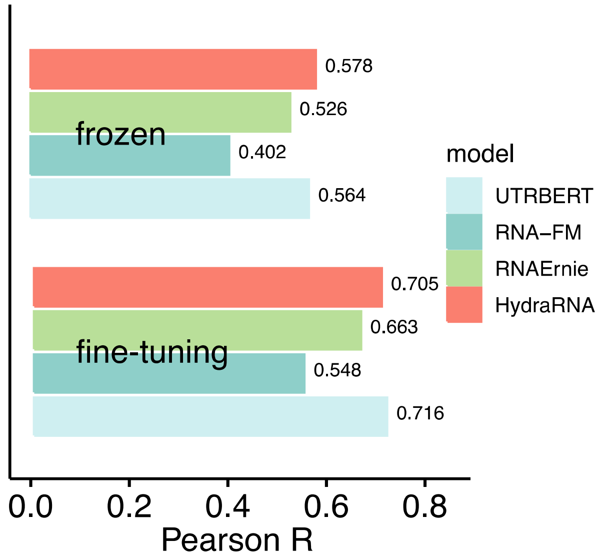


Figure S3. Performance comparison between models using frozen / all-parameters fine-tuning for the Jurkat dataset.

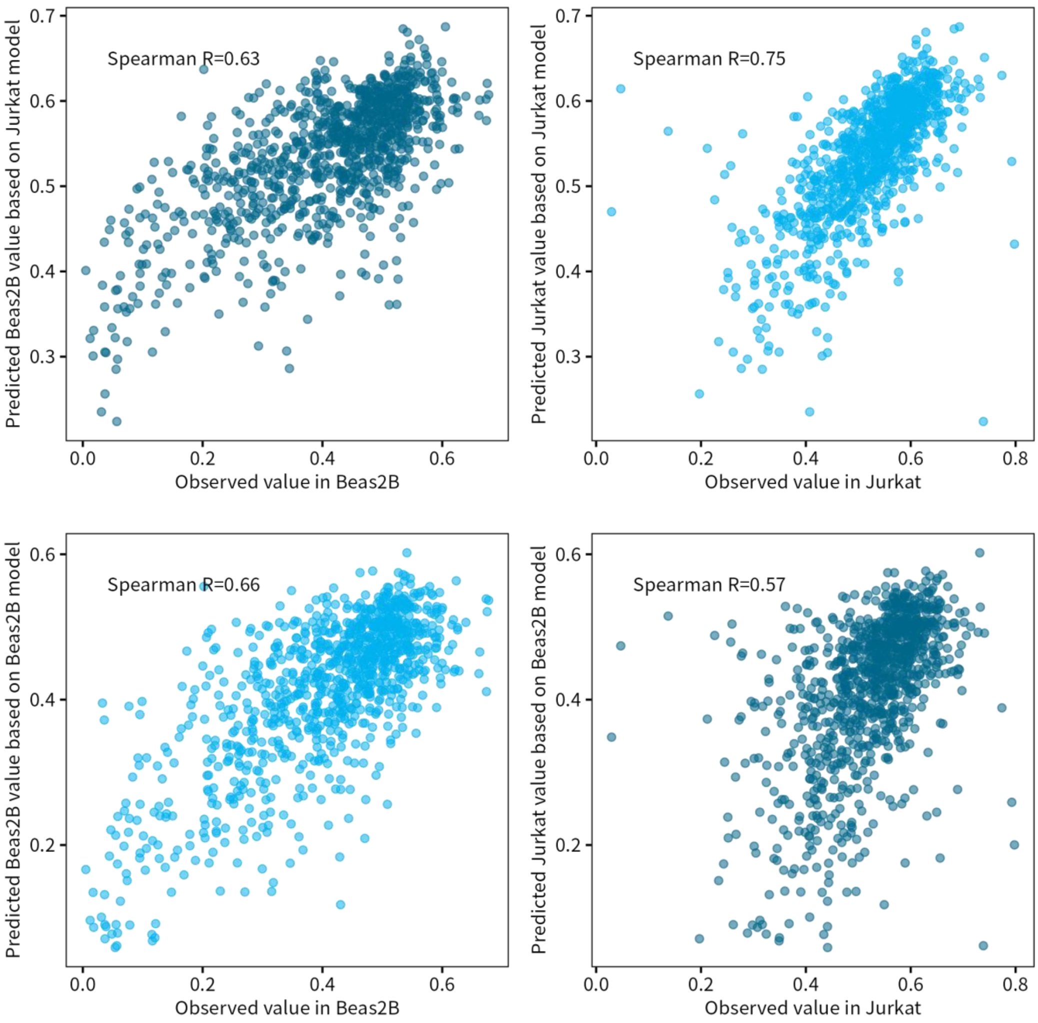


Figure S4. Performance of HydraRNA prediction within cell-line and across cell-lines.

Table S1. Comparison of pretraining process between different models.

| Model | HydraRNA | RNAErnie | RNA-FM |
|---|---|---|---|
| Data | RNAcentral , NCBI | RNAcentral | RNAcentral |
| #parameters | 84M | 105M | 86M |
| #sequences | 29M | 23M | 23M |
| length | 4096 nt | 512 nt | 1024 nt |
| GPU usage | 8 x 4090D (24GB) | 4 x V100 (32GB) | 8 x A100 (80GB) |
| training time | 92 hours | 250 hours | 30 days |
| estimated budget | 1472 RMB / 206 $ | 2000 RMB / 280 $ | 40320 RMB / 5645 $ |

Table S2. RBP Motif Data Obtained from mCrossBase

| RBP Name | Consensus | Score | Cell Line | RBP Group |
|---|---|---|---|---|
| hnRNPC | TTTTGGC | 4642.42 | HepG2 | high |
| TARDBP | TGTGTGA | 1305.02 | K562 | high |
| U2AF2 | TTCTTCC | 863.04 | K562 | high |
| QKI | ACTAACA | 682.69 | HepG2 | high |
| TIA1 | TTCTTTG | 479.58 | HepG2 | high |
| TAF15 | GTAGGGA | 402.58 | HepG2 | high |
| PUM2 | TGTACAG | 374.84 | K562 | high |
| IGF2BP2 | CCCCAGC | 157.45 | K562 | low |
| IGF2BP1 | CATTCCA | 133.34 | K562 | low |
| NSUN2 | CGTTCCT | 42.93 | K562 | low |
